# Enhanced Ca^2+^ Binding to EF-Hands through Phosphorylation of Conserved Serine Residues Activates MpRBOHB and Chitin-Triggered ROS Production

**DOI:** 10.1101/2023.10.05.559649

**Authors:** Takafumi Hashimoto, Kenji Hashimoto, Hiroki Shindo, Shoko Tsuboyama, Takuya Miyakawa, Masaru Tanokura, Kazuyuki Kuchitsu

**Affiliations:** Department of Applied Biological Science, Tokyo University of Science, Noda, Chiba, Japan; Graduate School of Biostudies, Kyoto University, Kitashirakawa-oiwakecho, Sakyo-ku, Kyoto, 606-8502, Japan; Department of Applied Biological Chemistry, Graduate School of Agricultural and Life Sciences, The University of Tokyo, 1-1-1 Yayoi, Bunkyo-ku, Tokyo, 113-8657, Japan

## Abstract

NADPH oxidases/RBOHs catalyze apoplastic ROS production and act as key signaling nodes, integrating multiple signal transduction pathways regulating plant development and stress responses. Although RBOHs have been suggested to be activated by Ca^2+^ binding and phosphorylation by various protein kinases, a mechanism linking Ca^2+^ binding and phosphorylation in the activity regulation remained elusive. Chitin-triggered ROS production required cytosolic Ca^2+^ elevation and Ca^2+^ binding to MpRBOHB in a liverwort *Marchantia polymorpha*. Heterologous expression analysis of truncated variants revealed that a segment of the N-terminal cytosolic region highly conserved among land plant RBOHs encompassing the two EF-hand motifs is essential for the activation of MpRBOHB. Within the conserved regulatory domain, we have identified two Ser residues whose phosphorylation is critical for the activation *in planta*. Isothermal titration calorimetry analyses revealed that phosphorylation of the two Ser residues increased the Ca^2+^ binding affinity of MpRBOHB, while Ca^2+^ binding is indispensable for the activation, even if the two Ser residues are phosphorylated. Our findings shed light on a mechanism through which phosphorylation potentiates the Ca^2+^-dependent activation of MpRBOHB, emphasizing the pivotal role of Ca^2+^ binding in mediating the Ca^2+^ and phosphorylation-driven activation of MpRBOHB, which is likely to represent a fundamental mechanism conserved among land plant RBOHs.

## 1. Introduction

Reactive oxygen species, frequently generated during fundamental cellular processes like aerobic respiration and photosynthesis, exhibit high reactivity and stand as primary contributors to damage resulting from a range of environmental stresses. Nonetheless, many eukaryotic organisms have evolved intricate enzymatic systems for the controlled production of ROS (Mittler, 2017). Notably, one such enzyme family, NADPH oxidases (Nox), plays a pivotal role in orchestrating this regulated ROS production (Sumimoto, 2008).

Nox exhibits a remarkable diversity in angiosperms, with *Arabidopsis thaliana* alone boasting ten distinct isozymes (Kaya et al., 2019). These isozymes participate in a myriad of vital biological processes integral to plant development and responses to environmental challenges (Kärkönen and Kuchitsu, 2015; Huang et al., 2019) such as polar tip growth of root hairs and pollen tubes (Takeda et al., 2008; Boisson-Dernier et al., 2013; Kaya et al., 2014; Lassig et al., 2014) as well as pathogen defense (Kadota et al., 2015).

In plant immunity, recognition of pathogen/microbes-associated molecular patterns (PAMPs/MAMPs), such as oligosaccharides derived from chitin, the major structural component of fungal cell wall, and peptides derived from pathogens, by specific pattern-recognition receptors (PRRs) triggers a cascade of defense responses (DeFalco and Zipfel, 2021). Early plasma membrane signaling events triggered by PAMPs such as chitin oligosaccharides and pathogen-derived signal molecules encompass a transient depolarization (Kuchitsu et al., 1993) and a range of ion fluxes (Kuchitsu et al., 1997) represented by transient cytosolic Ca^2+^ elevation (Blume et al., 2000; Lecourieux et al., 2002; Kadota et al., 2004; Kurusu et al., 2011; Segonzac et al., 2011; Hamada et al., 2012) and production of ROS including O_2_^-^, H_2_O_2_ and ^.^OH (Kuchitsu et al., 1995; Chinchilla et al., 2007), which depends on RBOH, Ca^2+^ and protein phosphorylation (Kuchitsu et al., 1995; Simon-Plas et al., 2002; Nühse et al., 2007; Zhang et al., 2007; Kadota et al., 2014).

ROS have been proposed to play multiple roles during the immune responses; including directly attacking the pathogens (Chen and Schopfer, 1999), inducing oxidative cross-linking as a substrate in the cell wall to invasion and spread of pathogens (Lamb and Dixon, 1997), reacting with lipids to produce reactive carbonyl species as signaling molecules (Biswas and Mano, 2021), and serving as signaling molecules (Bi et al., 2022; Dietz and Vogelsang, 2022).

Recent studies underscore the conservation of PAMP-triggered immune responses across land plants, extending to bryophytes. Chitin oligosaccharides trigger Ca^2+^ change in a moss *Physcomitrium (Physcomitrella) patens* (Galotto et al., 2020) and RBOH-mediated ROS production in a liverwort *Marchantia polymorpha* (Chu et al., 2023).

Plant NADPH oxidases/respiratory burst oxidase homologs (RBOHs) have N-terminal regulatory and C-terminal catalytic regions. An approximately 200 amino acids sequence domain containing two EF-hand motifs in the N-terminal regulatory region is well conserved in land plant RBOHs (Oda et al., 2010). Heterologous expression studies in HEK293T cells revealed that Ca^2+^ binding to the EF-hand motifs and phosphorylation synergistically activate all RBOHs of angiosperms and gymnosperms so far tested (Ogasawara et al., 2008; Takeda et al., 2008; Kimura et al., 2012; Takahashi et al., 2012; Kaya et al., 2014; Nickolov et al., 2022). In root hairs and pollen tubes, Ca^2+^ binding to the EF-hand motifs of RBOHs is required not only for the activation of the corresponding RBOHs but also for the proper polar tip growth, suggesting the importance of Ca^2+^ binding in the RBOH-mediated ROS production *in planta* (Takeda et al., 2008; Kaya et al., 2014).

Recent studies showed that RBOHs are activated by phosphorylation by various families of protein kinases such as Ca^2+^-dependent protein kinases (CPKs; Kobayashi et al. 2007; Dubiella et al. 2013), Ca^2+^-activated calcineurin B-like interacting protein kinases (CIPKs; Drerup et al. 2013; Kimura et al. 2013; Han et al. 2019), receptor-like cytoplasmic kinase (RLCKs; Kadota et al. 2014; Li et al. 2014, 2021), cysteine-rich receptor-like kinases (CRKs; Kimura et al. 2020), L-type lectin receptor-like kinases (Chen et al., 2017) and mitogen-activated protein kinase kinase kinase kinases (MAP4K) (Zhang et al., 2018a).

Overall, cumulative evidence suggests that plant RBOHs are activated by phosphorylation by various protein kinases and Ca^2+^ binding to the EF-hand motifs. However, the interrelationship between Ca^2+^ binding and phosphorylation in the activation of RBOHs has remained elusive. In the present study, we show that MpRBOHB, one of the two RBOHs in *Marchantia polymorpha*, is synergistically activated by Ca^2+^ and phosphorylation. Ca^2+^ binding to the EF-hand motifs is a requirement for its activation in a heterologous expression system using HEK293T cells as well as in chitin-triggered activation *in planta*. We have identified two Ser residues conserved in land plant RBOHs around the EF-hand motifs critical for the activation. Isothermal titration calorimetry analysis revealed that phosphorylation of the two Ser residues enhance the Ca^2+^-binding affinity. We propose this mechanism as a molecular basis of Ca^2+^- and phosphorylation-dependent synergistic activation of RBOHs, which may be conserved in land plants.

## 2. Materials and Methods

### 2.1 Plant materials and growth condition

*Marchantia polymorpha* Takaragaike-1 (Tak-1; male line) was cultured on half-strength Gamborg’s B5 medium (Gamborg et al., 1968) containing 1% sucrose and 1% agar under continuous white light (50 - 60 µmol m^−2^ s^−1^) at 21°C as described previously (Hashimoto et al., 2022).

### 2.2 Alignment of amino acid sequences

Amino acid sequences of RBOH homologs in land plants were collected from the database Phytozome (https://phytozome-next.jgi.doe.gov/) and Marpolbase (http://marchantia.info) and were aligned by the MAFFT program (Katoh et al., 2019) with FFT-NS-i strategy and default parameters.

### 2.3 Transgenic lines

Plant transformation mediated by *Agrobacterium tumefaciens* GV2260 was performed as described previously (Kubota et al., 2013; Tsuboyama et al., 2018). The binary vectors are constructed as follows.

The CRISPR/Cas9 genome editing system (Sugano et al., 2014) was employed to generate knockout lines. Oligonucleotides corresponding to designed guide RNAs (gRNA, listed in Table S1) were ligated into an entry vector pMpGE_En03 (Addgene #71535) and then transferred to a destination vector pMpGE010 (Addgene #71536).

To investigate the complementation of Mp*rbohB* knockout lines, we amplified a cDNA from the total RNA of Tak-1 plants by reverse transcription PCR and fused it to a promoter DNA by overlap extension PCR. The DNA fragment was cloned into the destination vector pMpGWB301 (Ishizaki et al., 2015) via pENTR/D-TOPO entry vector (Thermo Fisher Scientific). The variants of MpRBOHB examined in this study were generated by site-directed mutagenesis PCR and constructed into the binary vector as above. The DNA containing promotor region of Mp*RBOHB* was obtained by amplifying a 5-kb genomic DNA fragment upstream of the predicted initiation codon of Mp*RBOHB* gene by PCR and cloned into the pENTR/D-TOPO entry vector. All primers used in these experiments are listed in the supplemental table.

To visualize cytosolic Ca^2+^ level, we amplified the genetically encoded Ca^2+^ indicator GCaMP6f-mCherry (Waadt et al., 2017) from a plasmid kindly provided by Dr. Rainer Waadt, Muenster, Germany. The PCR-amplified gene was subcloned into pENTR/D-TOPO entry vector and transferred into pMpGWB303 (Ishizaki et al., 2015) via LR recombination. In the resultant T-DNA, the elongation factor 1α (EF1α) promoter drives the GCaMP6f-mCherry gene.

### 2.4 Plasmid construction

The cDNAs encoding MpRBOHB and its variants with 3 × FLAG tag at their 5’-end were cloned into the pcDNA3.1(-) vector (Invitrogen) for the quantitative measurement of ROS in HEK293T cells.

The cDNAs encoding MpRBOHB-(212-414), its phosphorylation mimic variant S223E/S406E (wild-type MpRBOHB^N^ and 2xSE MpRBOHB^N^, respectively), and linearized pRSF-2 Ek/LIC vector were amplified by PCR to add overlap sequences at their 5’- and 3’-end. All primers used in these experiments are listed in the supplemental table. The amplified cDNAs were cloned into pRSF-2 Ek/LIC vector by In-Fusion reaction with the linearized vector for isothermal titration microcalorimetry experiments. In-Fusion HD Cloning Kit (Clontech) was used for this reaction.

### 2.5 Measurement of cytosolic Ca^2+^ changes in plants

For Ca^2+^ imaging, GCaMP6f-mCherry-expressing lines were used. Gemmalings were grown on half-strength Gamborg’s B5 medium containing 0.1% sucrose and 1% agar under continuous white light. 2-day-old gemmalings were transferred to 6 well plates at five plants per well and further grown for 3 or 4 d. Before measurements, we added 3 mL/well of half-strength Gamborg’s B5 medium containing 0.1% sucrose to the plants in 6 well plates and left them at room temperature for over 1 h. For the pretreatment with a Ca^2+^ chelator *O*,*O*’-Bis(2-aminophenyl)ethyleneglycol-*N*,*N*,*N*’,*N*’-tetraacetic acid (BAPTA; Tokyo Chemical Industry), the plant materials were further kept at room temperature for more than 1 h. Fluorescence images were acquired by a fluorescence stereoscope (Nikon SMZ25) equipped with a camera (Nikon DS-Ri2), P2-SHR Plan Apochromatic 1X objective lens, and an excitation light source (Nikon INTENSILIGHT C-HGFIE). For fluorescence image acquisition, we used a 470/40 nm excitation filter, a 500 nm long-pass dichroic mirror and a 535/50 nm emission filter for GCaMP6f, and a 545/15 nm excitation filter, a 570 nm long-pass dichroic mirror, and a 620/30 nm emission filter for mCherry. *N*-acetylchitooligosaccharides (chitin fragments; Tokyo Chemical Industry, Product No. C2762) dissolved at a concentration of 4 g L^-1^ in the liquid medium with or without the inhibitor was added 5 min after the measurement of the background fluorescence. Image analysis was performed with ImageJ Fiji (Schindelin et al., 2012). The fluorescence intensity of the whole thallus of each plant was quantified.

### 2.6 Quantitative determination of ROS production *in planta*

For assays with Marchantia plants, gemmae were cultured in half-strength Gamborg’s B5 medium containing 0.1% sucrose under continuous white light for 6 d. Four gemmalings were placed in each well of a white 96 well plate and incubated in 120 μL of assay solution (Elix water containing 100 μM L-012 (Fuji Film Wako Pure Chemical Corporation) and 12.5 g L^−1^ horseradish peroxidase (HRP) (Fuji Film Wako Pure Chemical Corporation) for approximately 18 h at room temperature under darkness. Chitin treatment was performed by adding 13 μL of assay solution containing 10 g L^−1^ chitin oligosaccharides to each well (1 g L^−1^ final in wells). ROS production was determined by counting photons derived from L-012-mediated chemiluminescence using Luminometer Centro LB960 (Berthold Technologies).

### 2.7 Quantitative determination of the ROS-producing activity of MpRBOHB expressed in HEK293T cells

The ROS-producing activity of MpRBOHB heterologously expressed in HEK293T cells was examined as previously described (Ogasawara et al., 2008). Cells in each well of a white 96-well plate were transfected with 50 ng of pcDNA3.1(-) 3 × FLAG-MpRBOHB. ROS assay buffer was Hanks’ balanced salt solution (HBSS, Fuji Film Wako Pure Chemical Corporation, Product No. 085-09355) containing 250 μM L-012 and 4 mg L^−1^ horseradish peroxidase (Fuji Film Wako Pure Chemical Corporation). The expression of 3 × FLAG-tagged MpRBOHB proteins was confirmed by western blot analysis using an anti-3 × FLAG antibody (Sigma-Aldrich, Cat. No. F1804).

### 2.8 Estimation of intracellular Ca^2+^ concentration in HEK293T cells

HEK293T cells were cultured in a black 96 well plate under the same condition used in the ROS assay. Briefly, 1.0 × 10^5^ to 2.0 × 10^5^ cells mL^−1^ of cell suspensions were seeded into the 96 well plate and incubated for 3 d. Prior to the measurement, the media was aspirated from the cells and replaced with the Fura-2-loading solution (5 μM Fura-2-AM (Dojindo) dissolved in HBSS). After 30 min of incubation at 37°C, the loading solution was replaced with calcium solution (0 mM to 2 mM CaCl_2_ dissolved in HBSS). Fura-2 fluorescence was measured with an Infinite 200 Pro plate reader (Tecan). 1 μM ionomycin was added after measuring the baseline for 5 min. To determine the absolute Ca^2+^ concentration, Fura-2 fluorescence was calibrated with defined Ca^2+^ buffers generated with a Calcium Calibration Buffer Kit #1 (Invitrogen).

### 2.9 Preparation of recombinant proteins

Wild type and 2xSE variant of MpRBOHB^N^ with an N-terminal 6 × His tag were expressed in *Escherichia coli* Rosetta-gami 2(DE3) (Novagen). Cells were grown at 37°C in lysogeny broth (LB) medium containing 25 mg L^−1^ kanamycin. When the absorbance at 600 nm (OD_600_) of the cell culture reached 0.4, isopropyl-β-D(-)-thiogalactopyranoside (IPTG; Fuji Film Wako Pure Chemical Corporation) was added to a concentration of 1 mM to induce expression. Cells were grown for an additional 3 h at 37°C following IPTG induction and were harvested by centrifugation at 2000*g* for 10 min at 4°C. The harvested cells were disrupted by sonication. After centrifugation at 12000*g* for 30 min at 4°C, the soluble proteins were purified using Ni Sepharose 6 Fast Flow (Cytiva). 6 × His-tagged MpRBOHB^N^ proteins in each fraction were confirmed by western blot analysis using an anti-His tag antibody (MBL, Code No. PM032). The proteins were further purified by gel filtration chromatography, which were performed using an ÄKTA explorer 10S (Cytiva) and a Superdex HR 75 10/30 column (Cytiva) with the buffer containing 20 mM PIPES-KOH (pH 6.8) and 150 mM KCl. The purified proteins were concentrated to more than 100 µM using ultrafiltration filter units Vivaspin 6, 5,000 MWCO PES (Cytiva). The concentration was determined by absorbance at 280 nm with a molar extinction coefficient of 26930.

### 2.10 Isothermal titration microcalorimetry

Isothermal titration microcalorimetry (ITC) experiments were conducted at a controlled temperature of 25°C using an ITC instrument (MicroCal iTC200, Malvern Panalytical). The CaCl_2_ solution was prepared with the same buffer as MpRBOHB^N^ proteins (20 mM PIPES-KOH, pH 6.8, and 150 mM KCl). The sample cell was filled with 100 μM protein solution and titrated with 4.2 mM or 2.3 mM CaCl_2_ solution for wild-type MpRBOHB^N^ or 2xSE MpRBOHB^N^, respectively. For each titration, 19 consecutive 2 μL aliquots of the CaCl_2_ solution were injected at 120 s intervals.

Data analysis was performed using the Origin-ITC analysis package (Malvern Panalytical) in “One set of sites” mode. The appropriateness of the modes was evaluated by the χ^2^ value of curve-fitting to the ITC thermogram. Reaction enthalpies (Δ*H*), dissociation constants (*K*_d_), and stoichiometry (*N*) were determined directly from curve-fitting. The changes in the Gibbs free energy (Δ*G*) and entropy (Δ*S*) were determined by using the equation Δ*G* = −*RT*ln*K*_a_ = Δ*H* − *T*Δ*S*, where *R* and *T* are universal gas constant and absolute temperature.

## 3. Results

### 3.1 Ca^2+^-binding to the EF-hand motifs is critical for MpRBOHB activation

The deduced primary structure and domain structure of MpRBOHB (synonym of MpRBOH1 in Chu et al. (2023) and MpRBOH2 in Yotsui et al. (2023)) including the part of N-terminal regulatory region that contains two EF-hand motifs (Fig. 1A) as well as the C-terminal catalytic region highly conserved among eukaryotes (Sumimoto, 2008) was similar to those of RBOHs in angiosperms and gymnosperms (Nickolov et al., 2022).

**Fig. 1.**
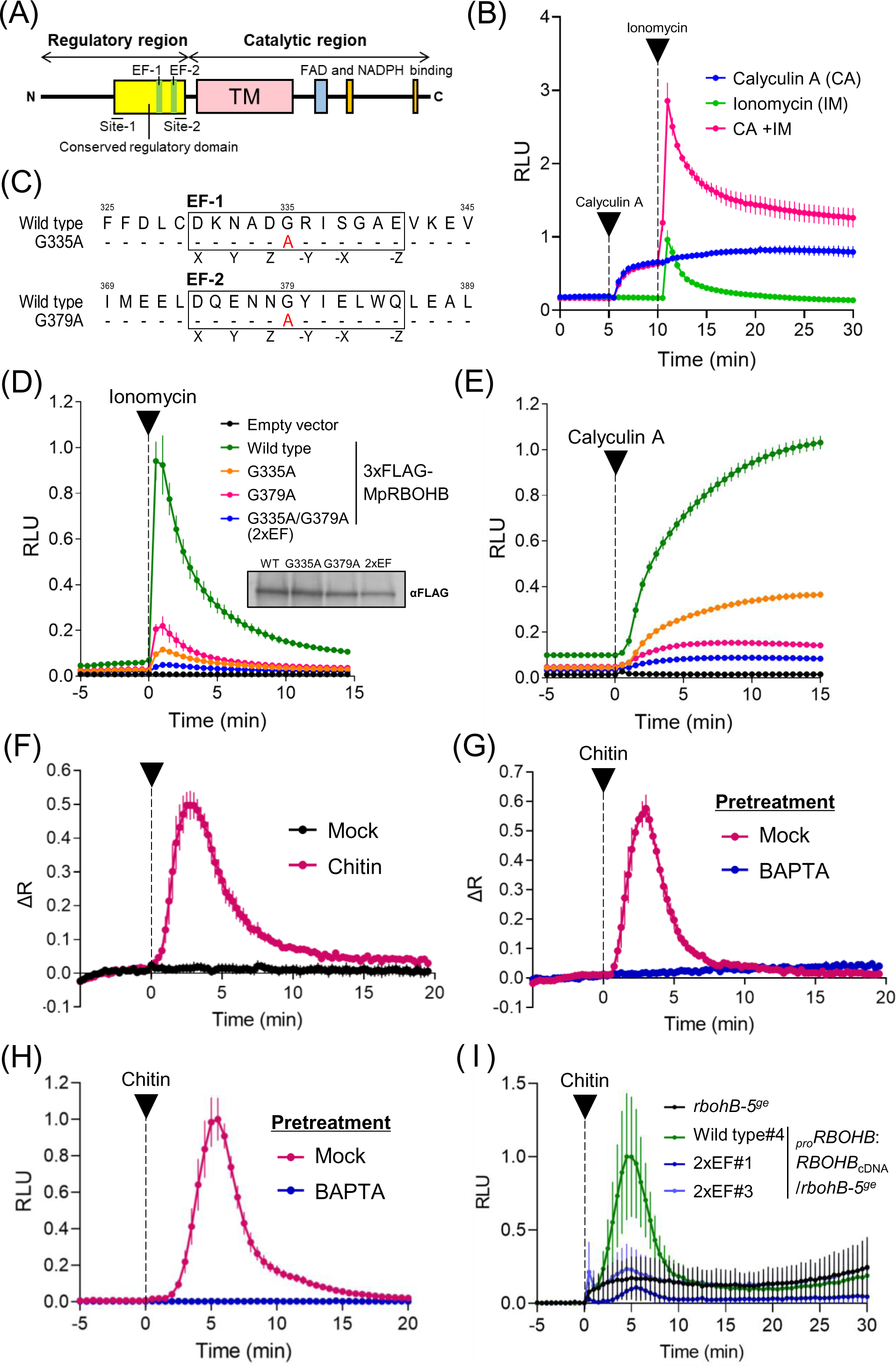
Ca^2+^ binding to the EF-hand motifs is required for activation of MpRBOHB. (A) A schematic diagram of MpRBOHB protein. The yellow box indicates the conserved domain in the N-terminal regulatory region, containing S223 and S406 (site-1, 2). (B) Synergistic activation of MpRBOHB by phosphorylation and Ca^2+^ was demonstrated using the HEK293T heterologous expression system. After baseline measurement, either 0.1 mM calyculin A (CA) or 1 mM ionomycin (IM), together with 1 mM CaCl_2_, was added at the time points indicated by the arrowheads. (C) Amino acid sequences of the EF-hand regions of MpRBOHB. Consensus amino acid positions involved in Ca^2+^-binding are indicated by X, Y, Z, -X, -Y, and -Z. Amino acid substitutions in the MpRBOHB variants tested in this study are shown below. (D) IM-induced activation of the MpRBOHB variants in HEK293T cells. Western analysis using an anti-FLAG tag antibody confirmed the expression of the MpRBOHB variants in the HEK293T cells. (E) CA-induced activation of the MpRBOHB variants in HEK293T cells. (F) 5-day- or 6-day-old gemmalings of Tak-1 expressing GCaMP6f-mCherry were treated with 0 g L^−1^ or 1 g L^−1^ chitin oligosaccharides. GCaMP6f and mCherry fluorescence was measured by a fluorescence stereomicroscope. ΔR = R – R_0_; R is GCaMP6f/mCherry ratio. (G) Treatments with 0 mM or 5 mM BAPTA were carried out 2 h prior to chitin treatment. (H) Chitin oligosaccharides-induced ROS production was measured by L-012 chemiluminescence in 7-day-old gemmalings of Tak-1. The chemiluminescence was expressed as relative luminescence units (RLU). Inhibitor treatments were carried out 1 h prior to chitin treatment. (I) Chitin oligosaccharides-induced ROS production in Mp*rbohB-5^ge^*and the complementation lines by wild type and G335A/G379A (2xEF) variant of MpRBOHB. Data are means ± SD for three replicates (B, D, E), five replicates (F, G), or six replicates (H, I).

To investigate the conservation of the synergistic activation by Ca^2+^ and phosphorylation observed in all angiosperm and gymnosperm species so far tested (Nickolov et al., 2022) in MpRBOHB, we heterologously expressed Mp*RBOHB* cDNA in HEK293T cells (Ogasawara et al., 2008). Cytosolic Ca^2+^ elevation introduced by the ionomycin (IM) treatment elicited transient activation of wild-type MpRBOHB (Fig. 1B). Treatment with a protein Ser/Thr phosphatase inhibitor calyculin A (CA), which is able to induce ROS production in rice cells (Kuchitsu et al., 1995) and phosphorylation of RBOH proteins in HEK293T cells (Ogasawara et al., 2008), triggered continuous activation. Pretreatment with CA enhanced IM-induced activation (Fig. 1B), indicating that basic activation mechanisms of MpRBOHB resembles homologs in other land plants, and MpRBOHB is synergistically activated by Ca^2+^ and phosphorylation.

IM-induced activation is mediated by Ca^2+^ binding to the EF-hand motifs in the N-terminal regulatory region (Ogasawara et al., 2008; Oda et al., 2010). The amino acid sequences of the EF-hand motifs are highly conserved in land plants, from bryophytes to angiosperms (Fig. S1).

We investigated the effect of Ca^2+^ binding to the EF-hand motifs on the activation of MpRBOHB. EF-hand motifs often share the consensus amino acid residues within the Ca^2+^ binding loop (Gifford et al., 2007). In general, substitution from negatively charged aspartate to uncharged asparagine at the first position X disrupts Ca^2+^ chelation. Glycine at the center facilitates the bend of the Ca^2+^ binding loop, and replacement to other amino acids (e.g., Ala) often interferes with chelation. Indeed, such mutations in each EF-hand motif of MpRBOHB (Figs 1C, S2A) severely suppressed the IM-induced and CA-induced activation in the HEK293T cells (Figs 1D, 1E, S2). Furthermore, the mutant MpRBOHB^G335A/G379A^ (2xEF), in which both of the two critical amino acids are replaced, showed almost no ROS-producing activity, significantly lower than the single amino acid-replaced mutants MpRBOHB^G335A^ and MpRBOHB^G379A^ (Figs 1D, E), suggesting that Ca^2+^ binding mediated by the two EF-hand motifs are crucial for the activation triggered not only by IM (Ca^2+^) but also CA (protein phosphorylation).

There are only two RBOH genes in *Marchantia polymorpha* (hereafter Marchantia), Mp*RBOHA* and Mp*RBOHB*. We generated knockout lines of Mp*RBOHA* and Mp*RBOHB* by genome editing (Fig. S3) and compared the chitin-triggered ROS production in independent lines. Chitin oligosaccharide triggered transient ROS production similarly in the wild type and in the mutants defective in *rbohA,* while no ROS production in both mutants defective in *rbohB,* indicating that MpRBOHB, but not MpRBOHA is activated upon recognition of chitin oligosaccharides (Fig. S4). These results are consistent with the recent report showing that chitin-triggered ROS production depend on MpRBOHB (Chu et al., 2023).

To evaluate the significance of Ca^2+^ in the chitin-triggered activation of MpRBOHB, we first examined whether chitin oligosaccharides also trigger the changes in cytosolic Ca^2+^ concentration ([Ca^2+^]_cyt_) in Marchantia. A fusion protein consisting of GCaMP6f, a Ca^2+^ sensor, and mCherry, a Ca^2+^-independent fluorescent protein (Waadt et al., 2017), was expressed in Tak-1 to monitor [Ca^2+^]_cyt_. Quantitative analyses of the fluorescence from GCaMP6f and mCherry with the fluorescence stereo microscope reveled that rapid and transient [Ca^2+^]_cyt_elevation was triggered by chitin treatment, but not by mock treatment (Fig. 1F).

The chitin-triggered Ca^2+^ elevation was suppressed by pre-treatment with a Ca^2+^chelator BAPTA (Fig. 1G), suggesting that chitin-triggered transient plasma membrane Ca^2+^ influx similarly to those observed in angiosperms is conserved in liverworts.

We next tested the effect of BAPTA pre-treatment on chitin-triggered ROS production. BAPTA pre-treatment drastically suppressed the chitin-triggered elevation of L-012 chemiluminescence (Fig. 1H). These data suggest that Ca^2+^ influx from the apoplast is necessary for chitin-triggered ROS production in Marchantia. To verify the role of direct Ca^2+^ binding to the EF-hand motifs in the Ca^2+^-mediated activation of MpRBOHB, we conducted a complementation analysis of the Mp*rbohB* knockout line with Mp*RBOHB* cDNA and its 2xEF variant. The wild-type MpRBOHB restored the chitin-triggered ROS production, but 2xEF MpRBOHB did not (Fig. 1I). These results suggest that Ca^2+^ binding to the EF-hand motifs is required for the activation of MpRBOHB *in planta* as well as in the heterologous expression system.

### 3.2 Two conserved phosphorylation sites critical for MpRBOHB activation and the effect of Ca^2+^ binding on the activation through these phosphorylation sites

The results in the heterologous expression system implied involvement of Ca^2+^ binding to the EF-hand motifs in phosphorylation-dependent regulation (Fig. 1E). To investigate the role of Ca^2+^ binding in the phosphorylation-dependent regulation among the conserved regulatory mechanisms, we first explored the Ser/Thr residues, which are critical for the activation and common in land plant RBOHs.

Putative phosphorylation sites of RBOHs have been reported in both the variable domain and conserved domain of the N-terminal regulatory region. We firstly tested the significance of the conserved regulatory domain using the heterologous expression system by analyzing several truncated variants of MpRBOHB (Fig. 2A). The variable domain lacking variant, MpRBOHB^Δ1-211^, exhibited activations by CA and IM just like wild-type MpRBOHB. However, the variant lacking both the variable and conserved domain, MpRBOHB^Δ1-421^, did not respond to CA as well as IM (Fig. 2A). These results suggest that the conserved regulatory domain encompassing the two EF-hand motifs is necessary and sufficient for the common activation mechanism that relies on both Ca^2+^ binding and phosphorylation.

**Fig. 2.**
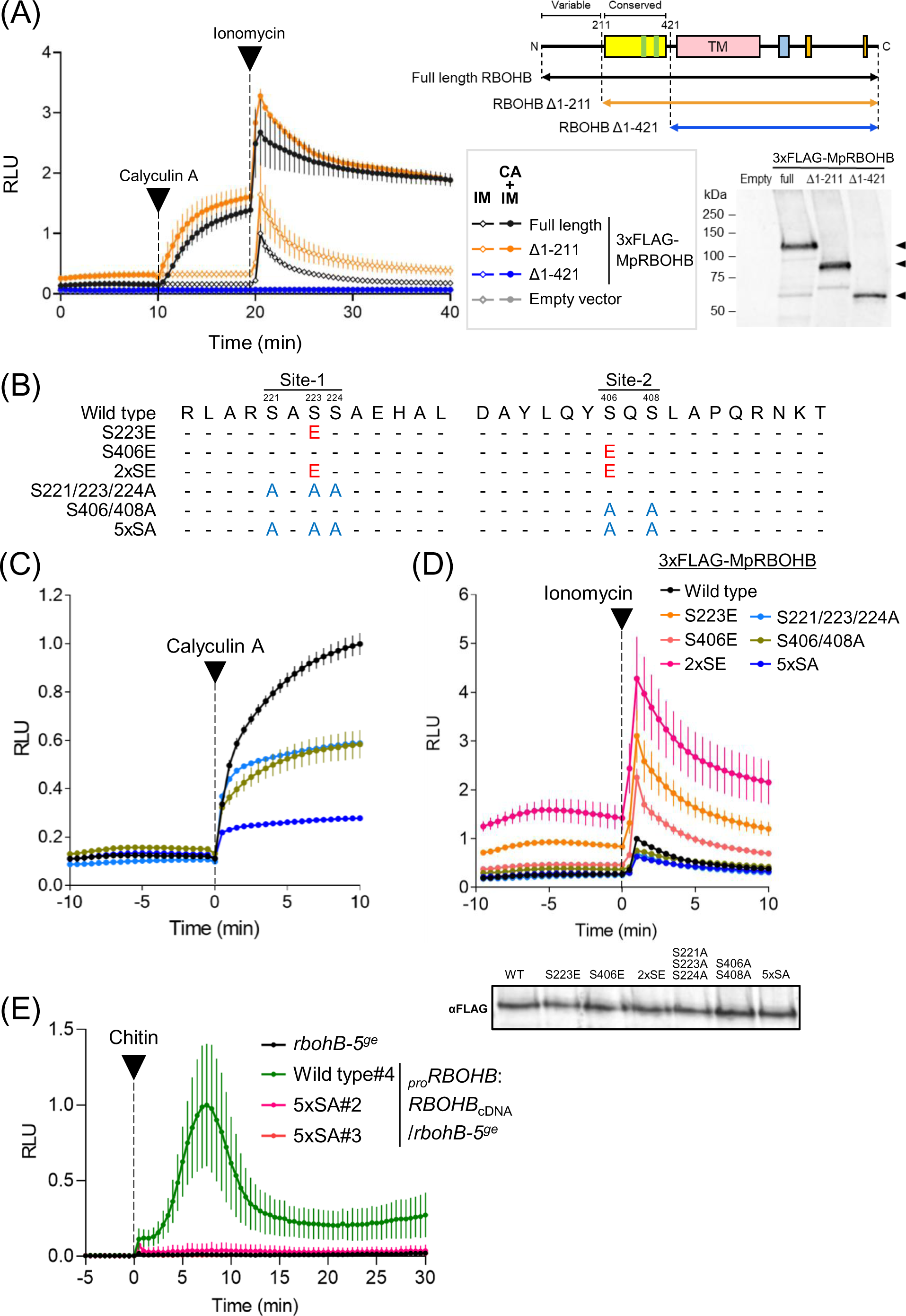
Two Ser residues in the conserved regulatory domain are involved in the activation of MpRBOHB (A) The activation of the truncated MpRBOHB by calyculin A and ionomycin was demonstrated using the HEK293T expression system. The truncated variants tested are depicted below the schematic structure of MpRBOHB protein. Western blot is shown in lower right panel, and arrowheads indicate the truncated MpRBOHB variants. Data are means ± SD for three replicates. (B) Amino acid sequences of the phosphorylation sites of MpRBOHB. Amino acid substitutions in the phosphorylation-mimic and phosphorylation-dead variants tested in this study are shown below. (C) Calyculin A-induced activation and (D) Ionomycin-induced activation of MpRBOHB variants expressed in HEK293T cells. Western analysis using an anti-FLAG tag antibody confirmed the expression of the MpRBOHB variants in the HEK293T cells. Data are means ± SD for three replicates. (E) Chitin oligosaccharides-induced ROS production in Mp*rbohB-5^ge^* and the complemented lines by wild type and 5xSA variant of MpRBOHB. Data are means ± SD for six replicates.

We conducted an alignment of amino acid sequences within the conserved regulatory domain and found that S223 and S406 are widely conserved among land plant RBOHs (Fig. S5). Remarkably, these residues correspond to S163 and S347 in Arabidopsis AtRBOHD, both of which have been reported to undergo phosphorylation *in planta* (Kadota et al., 2014). To investigate the impact of phosphorylation on the activation of MpRBOHB, we introduced substitutions at S223 and S406 with Ala to inhibit their phosphorylation. To avoid neighboring phosphorylation, we also substituted adjacent Ser residues, namely S221, S224, and S408 with Ala (Fig. 2B). Notably, these variant constructs, particularly the 5xSA mutant, exhibited reduced activation upon treatment with CA (Fig. 2C), suggesting that these Ser residues play a crucial role in the phosphorylation-triggered activation of MpRBOHB.

Given that CA treatment enhanced IM-induced activation (Fig. 1B), we further substituted S223 and S406 with Glu to mimic their phosphorylated state (S223E and S406E; Fig. 2B). Subsequent examination of their IM-induced activation revealed that the S223E and S406E variants displayed higher activity compared to wild-type MpRBOHB, both in the steady state and IM-induced activation. Moreover, the double phosphorylation-mimic variant involving the two Ser residues (2xSE; Fig. 2B) exhibited significantly greater activity than the S223E and S406E variants (Fig. 2D). Conversely, the 5xSA variant displayed lower IM-induced activity than wild-type MpRBOHB (Fig. 2D). These results collectively suggest that phosphorylation of S223 and S406 is significant in enhancing Ca^2+^-triggered activation in MpRBOHB.

To assess the functional consequences of these two Ser residues *in planta*, we introduced the 5xSA variant of MpRBOHB cDNA into the Mp*rbohB* knockout background of Marchantia. In contrast to wild-type MpRBOHB, the 5xSA MpRBOHB failed to restore chitin-triggered ROS production (Fig. 2E), underscoring the critical role of phosphorylation at S223 and S406 in chitin-triggered ROS production.

We proceeded to investigate the influence of Ca^2+^ binding to the EF-hand motifs on the regulation of the two phosphorylated Ser residues. In comparison to wild-type MpRBOHB, the 2xSE MpRBOHB variant exhibited elevated basal activity and increased activation in response to IM stimulation. However, MpRBOHB^S223E/G335A/G379A/S406E^ (2xSE/2xEF MpRBOHB) showed no difference from 2xEF MpRBOHB, not only at IM-induced activation but also at basal activity (Fig. 3A). These results suggest that the regulation by phosphorylation of the two Ser residues depends on Ca^2+^ binding in the heterologous expression system. To explore the dependency of these interactions *in planta*, we expressed the 2xSE and 2xSE/2xEF variants of Mp*RBOHB* cDNA in the Mp*rbohB* knockout background of Marchantia. Complementation lines expressing 2xSE MpRBOHB exhibited chitin-triggered ROS production, which appeared more sustained compared to the wild-type MpRBOHB complementation line. Conversely, the 2xSE/2xEF MpRBOHB complementation lines did not display significant ROS production in comparison to the Mp*rbohB* knockout line (Fig. 3B, C). These findings, both from the heterologous expression system and *in planta*, collectively underscore that phosphorylation of the two Ser residues alone is insufficient to activate MpRBOHB; Ca^2+^ binding is an indispensable requisite for activation.

**Fig. 3.**
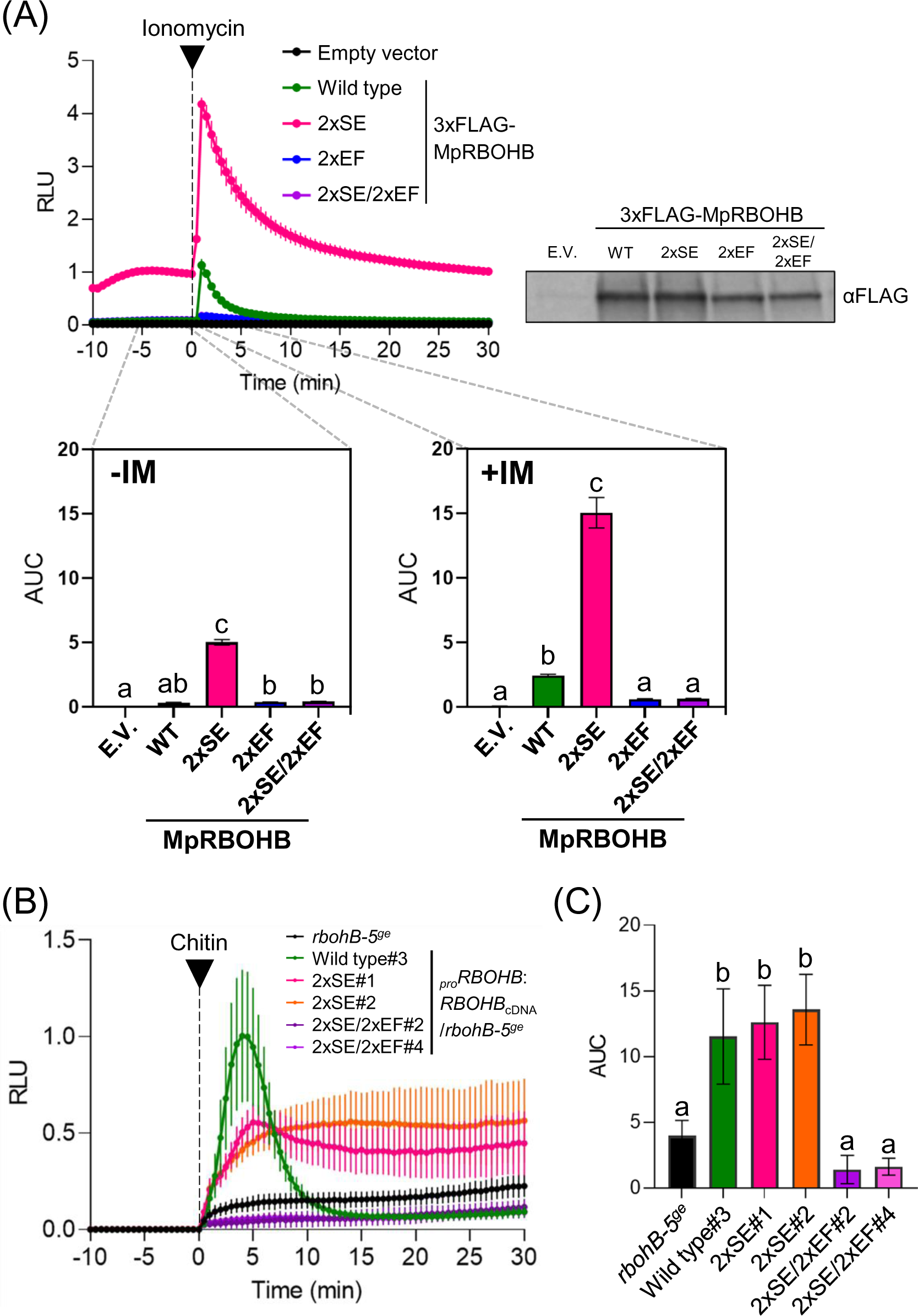
Ca^2+^ binding to the EF-hand motifs is required for activation of MpRBOHB even when S223 and S406 are phosphorylated both in HEK293T cells and *in planta* (A) ROS-producing activity of MpRBOHB variants in HEK293T cells. Area under curve values for 5 min before and after ionomycin treatment (-IM and +IM) are shown. Data are means ± SD for three replicates. Western analysis using an anti-FLAG tag antibody confirmed the expression of the MpRBOHB variants in the HEK293T cells. (B) Chitin oligosaccharides-induced ROS production in Mp*rbohB-5^ge^*and the complemented lines by wild type and 2xSE and 2xSE/2xEF variants of MpRBOHB. Data are means ± SD for six replicates. (C) Area under curve values for 15 min after chitin oligosaccharides treatment in (B). Equal letters above bars indicate p > 0.05, Tukey’s multiple comparison test.

### 3.3 Phosphorylation of the conserved two Ser residues increases the Ca^2+^ binding affinity of the N-terminal regulatory region of MpRBOHB

In the heterologous expression system, 2xSE MpRBOHB showed more robust IM-induced activation than wild-type MpRBOHB, like after CA treatment (Fig. 2D). We supposed that phosphorylation of the two conserved Ser residues enhances sensitivity to Ca^2+^, resulting in the increment of IM-induced activity of MpRBOHB. To examine this possibility, we compared the Ca^2+^ concentration dependency of the activity in wild-type MpRBOHB and 2xSE MpRBOHB using the heterologous expression system. Intracellular Ca^2+^ concentration was estimated by measuring fluorescence of an intracellular Ca^2+^ indicator Fura 2-AM (Fig. S6). At high Ca^2+^ concentration, both wild-type and 2xSE MpRBOHB showed IM-induced activation in a Ca^2+^ concentration-dependent manner. However, at low Ca^2+^ concentration, 2xSE MpRBOHB is activated in a Ca^2+^ concentration-dependent manner even though wild-type MpRBOHB did not show Ca^2+^ concentration dependency (Fig. 4). These results suggest that phosphorylation of S223 and S406 increases Ca^2+^ sensitivity of MpRBOHB.

**Fig. 4.**
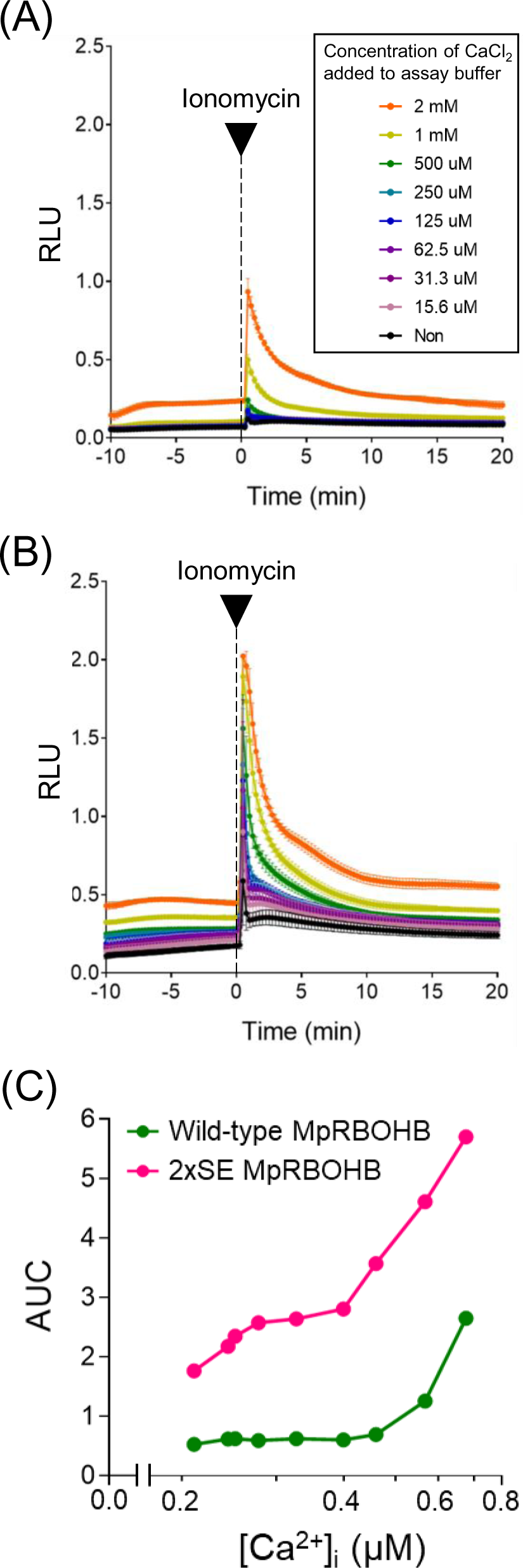
Ca^2+^ concentration dependency of ionomycin-triggered activity of wild-type and phosphorylation-mimic MpRBOHB Ionomycin-induced activation of wild-type (A) and 2xSE (B) MpRBOHB in HEK293T cells under indicated Ca^2+^ conditions. Data are means ± SD for three replicates. Data are means ± SD for three replicates. (C) Area under curve values for 5 min after ionomycin treatment in (A) and (B) were plotted against the estimated intracellular Ca^2+^ concentration ([Ca^2+^]_i_) in HEK293T cells.

In our pursuit to validate the hypothesis that phosphorylation of the two Ser residues leads to an enhancement of Ca^2+^ binding, thereby increasing Ca^2+^ sensitivity, we conducted a comparative examination of the Ca^2+^-binding affinity between wild-type and 2xSE MpRBOHB utilizing isothermal titration calorimetry (ITC). To facilitate these experiments, we generated the N-terminal domain of MpRBOHB (MpRBOHB^N^), which encompasses S223, S406, and two EF-hand motifs, as a recombinant protein fused with a 6 × His-tag (Figs 5A, S7). Both wild-type MpRBOHB^N^ (Fig. 5B) and 2xSE MpRBOHB^N^ (Fig. 5C) showed endothermic reactions associated with CaCl_2_ titration, and the parameters calculated from the thermograms are summarized in Fig. 5D.

**Fig. 5.**
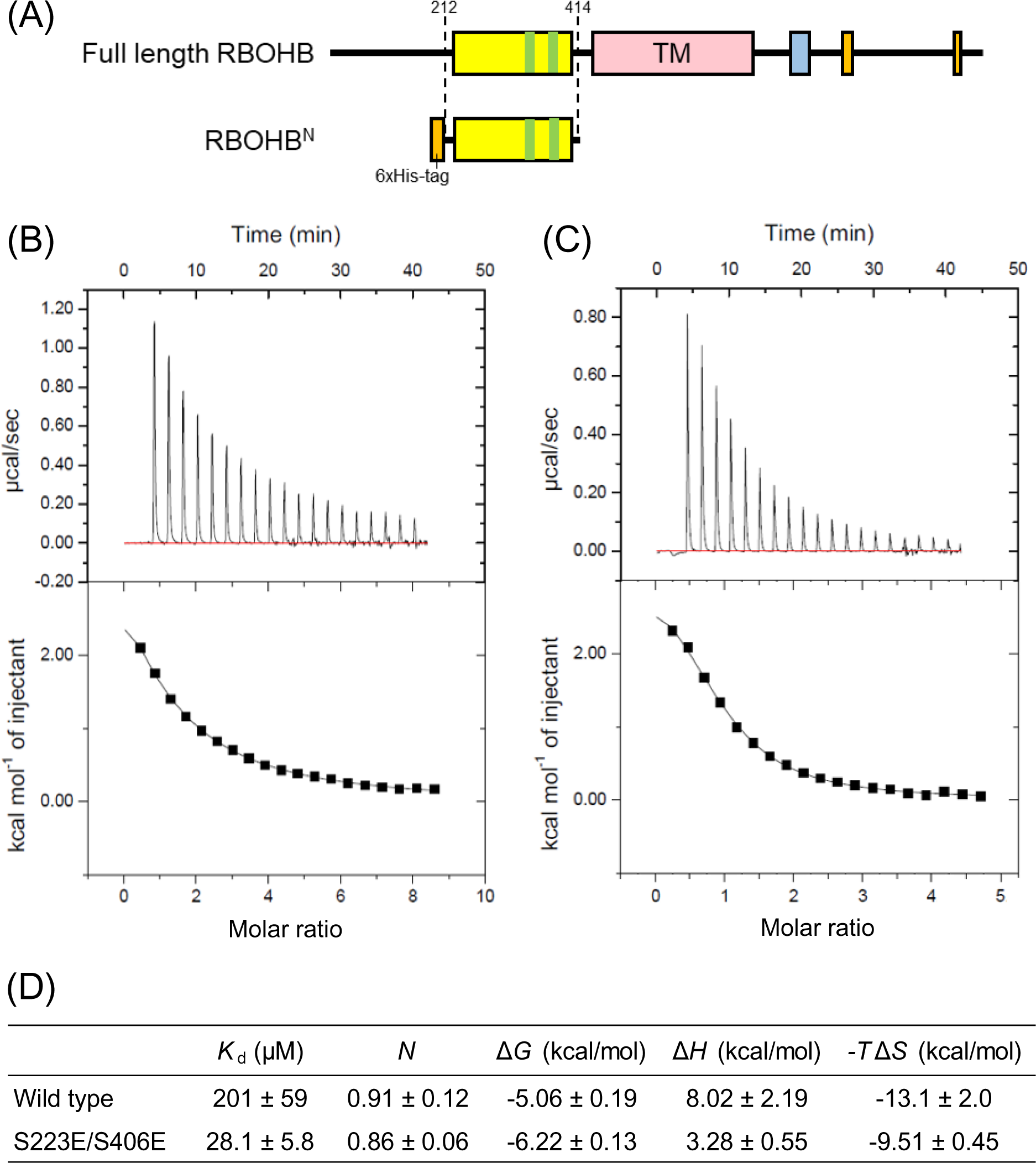
Microcalorimetric titration of conserved regulatory domain of wild-type and phosphorylation-mimic MpRBOHB with CaCl_2_ solution (A) A schematic diagram of MpRBOHB^N^ protein. The trace of the titration obtained by 19 injections of (B) 4.2 mM CaCl_2_ into 100 μM wild-type MpRBOHB^N^or (C) 2.3 mM CaCl_2_ into 100 μM 2xSE MpRBOHB^N^. In the lower panels, integrated heats for each injection versus the molar ratio of MpRBOHB^N^ to CaCl_2_are shown. The solid line indicates the curve fit of the data using the One Set of Sites model. These are representative data from experiments performed in triplicate. Similar results were obtained in the other two independent replicate experiments. (D) Thermodynamic parameters of wild type and phosphorylation mimicked (S223E/S406E) MpRBOHB^N^ obtained by ITC measurements. *H*: reaction enthalpy, *K*_d_: dissociation constant, *N*: stoichiometry, *G*: the Gibbs free energy, *S*: entropy. Data are means ± SD for three replicates.

The *N* values suggested that MpRBOHB^N^ binds Ca^2+^ at a single site, despite the presence of two EF-hand motifs, which aligns with the previous structural analysis of OsRBOHB (Oda et al., 2010). The mean of the *K*_d_ values for Ca^2+^ affinity, determined from three independent measurements using separately prepared samples was 201 μM for the wild-type MpRBOHB^N^ and 28.1 μM for the 2xSE MpRBOHB^N^, clearly demonstrating that the 2xSE variant exhibits a significantly higher Ca^2+^-binding affinity compared to the wild type (Fig. 5D). These results suggest that phosphorylation of S223 and S406 increases the Ca^2+^-binding affinity of the conserved N-terminal regulatory domain of MpRBOHB. This discovery provides compelling evidence supporting the hypothesis that phosphorylation indeed augments the Ca^2+^ binding capacity, shedding light on the mechanistic basis for the heightened Ca^2+^ sensitivity observed in the 2xSE variant.

## 4. Discussion

### 4.1 Interrelationships between Ca^2+^ binding and phosphorylation in the regulation of RBOH

ROS play crucial roles in regulating various biological processes critical for plant development, growth, and adaptation to biotic and abiotic stresses (Kärkönen and Kuchitsu, 2015; Mittler et al., 2022). Among the ROS-producing plant enzymes, RBOHs act as central signaling hubs, integrating multiple signal transduction pathways. ROS production by RBOHs is contingent not only on the quantitative regulation of RBOH protein, including transcriptional control of the *RBOH* genes and degradation of RBOH proteins but also on the precise control of the ROS-producing activity of RBOHs (Lee et al., 2020).

The activation of RBOHs involves the dual regulatory factors of Ca^2+^ binding and phosphorylation by various protein kinases (Kobayashi et al., 2007; Takeda et al., 2008; Drerup et al., 2013; Dubiella et al., 2013; Kimura et al., 2013, 2020; Kaya et al., 2014; Li et al., 2014, 2021; Zhang et al., 2018a; Han et al., 2019). While Ca^2+^ and phosphorylation have been suggested to cause synergistic activation of RBOHs in angiosperm and gymnosperm species as an output (Ogasawara et al., 2008; Takeda et al., 2008; Kimura et al., 2012; Takahashi et al., 2012; Kaya et al., 2014; Nickolov et al., 2022), the roles of these regulatory factors were largely unclear. In this study, we demonstrated that phosphorylation of two highly conserved Ser residues, S223 and S406, enhances the binding affinity and sensitivity of MpRBOHB to Ca^2+^ (Figs 4, 5) to play a pivotal role in the activation of MpRBOHB (Figs 2C-E). Notably, we also showed that even when MpRBOHB is phosphorylated, Ca^2+^ binding mediated by the EF-hand motifs remains an indispensable requisite for activation (Figs 1E, 1I, 3A-C). These findings support the notion that heightened Ca^2+^ affinity resulting from the phosphorylation of the two Ser residues enhances the Ca^2+^-triggered activation. Furthermore, our truncation analysis highlights the necessity of the conserved regulatory domain containing the two Ser residues and the two EF-hand motifs for Ca^2+^- and phosphorylation-mediated MpRBOHB activation (Fig. 2A). In Arabidopsis, S163 and S347 of AtRBOHD, corresponding to S223 and S406 of MpRBOHB (Fig. S5), are phosphorylated in response to chitin-, flg22-, and elf18 (Kadota et al., 2014). It is also suggested that the corresponding residues in AtRBOHC and AtRBOHF are phosphorylated by CBL1-CIPK26 and OST1, respectively (Sirichandra et al., 2009; Zhang et al., 2018b). In Marchantia, we found that phosphorylation of S223 and S406 can individually affect MpRBOHB activity (Figs 2C, D). S406 has recently been shown to be phosphorylated by MpPBLa in response to chitin (Chu et al., 2023). Collectively, we propose that the increase in Ca^2+^ affinity resulting from the phosphorylation of these Ser residues may represent a conserved regulatory mechanism across other land plant RBOHs. We posit that these mechanisms, orchestrated by the conserved regulatory domain, constitute fundamental regulatory processes governing land plant RBOHs.

MpRBOHB-mediated ROS production required the binding of Ca^2+^ to the EF-hand motifs in HEK293T cells (Fig. 1D), and similarly, chitin-triggered ROS production required Ca^2+^ binding to the EF-hand motifs *in planta* (Fig. 1I). This constitutes the first direct evidence showcasing the necessity of Ca^2+^ binding to RBOH for ROS production *in planta,* implying that RBOHs serves as a direct sensor of Ca^2+^ in the vicinity of plasma membrane *in planta* to initiate ROS production.

While earlier studies have proposed that Ca^2+^ binding to EF-hand motifs is not necessary for calyculin A (CA)-induced activation of Arabidopsis and rice RBOHs (Ogasawara et al., 2008; Kimura et al., 2012; Takahashi et al., 2012), these observations do not necessarily indicate fundamental differences in the regulatory mechanisms between liverwort and angiosperms. We showed that CA- and IM-induced activation, along with their synergistic effect, is conserved in MpRBOHB, mirroring the responses observed in seed plants (Fig. 1B). The C-terminal catalytic region and the approximately 200 amino acids sequence domain containing two Ca^2+^-binding EF-hand motifs in the N-terminal regulatory region are well conserved in land plant RBOHs (Oda et al., 2010). Moreover, a single EF-hand mutant of OsRBOHB exhibited partially suppressed CA-induced activation (Takahashi et al., 2012), implying the involvement of Ca^2+^ binding to EF-hand motifs in this suppression. The partial suppression of CA-induced activation was also shown in single EF-hand mutants G335A and G379A of MpRBOHB, and further suppression was demonstrated in the 2xEF mutant (Fig. 1E), indicating the participation of both EF-hand motifs in CA-induced activation. Consequently, we posit that Ca^2+^ binding-dependent regulation is a shared feature among land plant RBOHs.

Nevertheless, minor differences in amino acid sequences are present in the regulatory and catalytic regions of land plant RBOHs. For example, AtRBOHD possesses a phosphorylation site not conserved in MpRBOHB and is a target of CRK2 in the flg22 response (Kimura et al., 2020). Thus, in some molecular species of RBOHs, several mechanisms acquired during evolution might enable Ca^2+^-independent activation.

The apparent Ca^2+^ affinities of the truncated proteins used for the ITC experiments are low compared to physiological [Ca^2+^]_cyt_, even in the phosphorylation-mimic variant with elevated Ca^2+^-binding affinity (Fig. 5D). We postulate several plausible explanations for this apparent discrepancy. Firstly, for the ITC measurements, we had to use the truncated proteins instead of the whole RBOH protein. In rice OsRBOHB, it is suggested that the N-terminal regulatory region interacts with the C-terminal catalytic region in a Ca^2+^-independent manner (Oda et al., 2010). TnC, the Ca^2+^-binding subunit of troponin, can bind to Ca^2+^ independently, and its complex with TnI exhibits higher Ca^2+^ affinity than free TnC (Grabarek et al., 1986). From these observations, it is conceivable that the low Ca^2+^ affinity in our ITC experiments may be due to the absence of interaction with the catalytic region. Secondly, RBOH proteins predominantly localize to the plasma membrane, particularly within the plasma membrane microdomains, together with some proteins involved in RBOH regulation (Nagano et al., 2016), and Ca^2+^ permeable channels might participate in it (Shabala et al., 2015). When plasma membrane Ca^2+^ channels are activated, prompting an influx of Ca^2+^, the local concentration of Ca^2+^ in the vicinity of these microdomains is anticipated to surpass the average Ca^2+^ concentration, at least temporarily. In this context, RBOH proteins may function as Ca^2+^ sensor proteins, converting the Ca^2+^ signal initiated by plasma membrane Ca^2+^ channels into a ROS signal. In this scenario, the Ca^2+^-dependent phosphorylation of RBOHs by Ca^2+^-activated protein kinases might also work synergistically to enhance ROS production. Irrespective of the underlying mechanism, our findings unequivocally demonstrate that the Ca^2+^ affinity of the conserved N-terminal domain of the phosphorylation-mimic variant was higher than that of the wild type.

Although ITC experiments suggest that only one of the two EF-hand motifs binds to Ca^2+^ (Fig. 5D), both the G335A variant and G379A variant of MpRBOHB exhibited slight Ca^2+^-triggered activation in HEK293T cells. This observation implies that both EF-hand motifs can bind to Ca^2+^ in HEK293T cells (Fig. 1D). The Ca^2+^ binding ability of the two EF-hand motifs *in vivo* is worth investigating.

### 4.2 Roles of Ca^2+^ in the regulation of ROS signaling during immune responses

Ca^2+^ is a well-known second messenger in responses to biotic and abiotic stimuli (Kudla et al., 2010). During immune responses, a variety of Ca^2+^ sensor proteins can decode Ca^2+^-influx signatures (Lu and Tsuda, 2021). Ca^2+^ influx is required for ROS production triggered by PAMPs/MAMPs, including chitin oligosaccharides (Kadota et al., 2004; Kurusu et al., 2011; Segonzac et al., 2011). We demonstrated a rapid and transient [Ca^2+^]_cyt_ elevation induced by PAMP in Marchantia (Fig. 1F). To our knowledge, this is the first report of stimulus-triggered [Ca^2+^]_cyt_ elevation in Marchantia, and the temporal pattern of [Ca^2+^]_cyt_ elevation resembles that of angiosperm leaves and cultured cells. We also showed that [Ca^2+^]_cyt_ elevation is required for PAMP-triggered ROS production in Marchantia (Figs 1G, 1H). These observations suggest that Ca^2+^-induced ROS production in plant immunity is a shared mechanism among land plants, from bryophytes to angiosperms.

Cell wall peroxidases have also been reported to play a role in ROS accumulation in response to various elicitors (Daudi et al., 2012). However, rapid ROS production triggered by PAMPs, including chitin, relies on RBOHs (Simon-Plas et al., 2002; Kobayashi et al., 2007; Nühse et al., 2007; Zhang et al., 2007; Kadota et al., 2014). Mp*rbohB* knockout lines exhibited almost complete impairment in chitin-triggered ROS production (Fig. S4; Chu et al. 2023), indicating the major role of RBOH in the chitin-triggered ROS production in Marchantia, similarly to angiosperms. The Ca^2+^-dependent activation of RBOHs seems to play a fundamental role in the ROS production in pattern-triggered immunity.

In Marchantia plants, the lack of chitin-triggered activation of the phosphorylation-dead variant of MpRBOHB (Fig. 2E) may be attributed to an insufficient increase in Ca^2+^ affinity through phosphorylation to respond adequately to the elevated [Ca^2+^]_cyt_. On the other hand, the complementation lines expressing the phosphorylation-mimic MpRBOHB exhibit continuous chitin-triggered ROS production (Fig. 3B), possibly due to their higher Ca^2+^ affinity compared to wild-type MpRBOHB.

### 4.3 Complex regulatory mechanisms of RBOHs by Ca^2+^, protein kinases, and other regulatory factors

Plant RBOHs are involved in numerous physiological processes, such as polar tip growth (Foreman et al., 2003; Boisson-Dernier et al., 2013; Kaya et al., 2014; Lassig et al., 2014; Nestler et al., 2014; Wang et al., 2018), primary and lateral root development (Jiao et al., 2013; Li et al., 2015), seed germination (Müller et al., 2009; Li et al., 2017), salt tolerance (Ma et al., 2012; Kurusu et al., 2015), and wound responses (Sagi et al., 2004; Kumar et al., 2007; Miller et al., 2009), cold (Piotrovskii et al., 2011; Kawarazaki et al., 2013; Zhang et al., 2018c) and drought stress (Jiang and Zhang, 2002; Wang et al., 2016) in close connection with Ca^2+^ signaling (Dodd et al., 2010; Kurusu et al., 2013). Given the conservation of EF-hand motifs and Ser/Thr residues corresponding to S223 and S406 of MpRBOHB, the regulatory mechanisms proposed in this study might contribute to activation of various RBOHs in diverse physiological functions. Indeed, the requirement of Ca^2+^ binding and phosphorylation of the residue highlighted in this study for RBOH function is indicated in AtRBOHC during root hair tip growth (Takeda et al., 2008). On the other hand, besides Ca^2+^and protein kinases, various other post-translational modifications, such as phosphatidic acid (Zhang et al., 2009), nitrosylation (Yun et al., 2011) and persulfidation (Shen et al., 2020), and binding proteins, such as small GTPase Rac/ROP (Kawasaki et al., 1999; Ono et al., 2001; Wong et al., 2008) and a low temperature-inducible protein AtSRC2 (Kawarazaki et al., 2013), have been proposed to regulate RBOHs in several physiological functions. Even though enhancement of Ca^2+^ binding to the EF-hand motifs triggered by the phosphorylation of the two conserved Ser residues proposed in the present study seems to the fundamental conserved mechanism among land plant RBOHs, interplay with other posttranslational modification of RBOHs as well as interaction with other RBOH-binding proteins require future studies. Considering the limited genetic redundancy of RBOHs and related regulatory factors, *Marchantia polymorpha* appears to be a promising model organism for comprehensively studying the broader landscape of RBOH regulation and ROS signaling.

## Author contributions

Conception and design: Takafumi Hashimoto, Kenji Hashimoto, Kazuyuki Kuchitsu. Data collection and analysis: Takafumi Hashimoto, Kenji Hashimoto, Hiroki Shindo, Shoko Tsuboyama, and Takuya Miyakawa. Writing the manuscript: Takafumi Hashimoto, Kazuyuki Kuchitsu, Kenji Hashimoto, and Takuya Miyakawa. Providing resources: Kazuyuki Kuchitsu, Masaru Tanokura, and Takuya Miyakawa. All authors read and approved the final manuscript.

## Acknowledgements

This work was supported in part by JSPS KAKENHI Grant Numbers, JP26111008, JP20H02990, and JP22H04734. We thank Dr. Hirofumi Nakagami (Max Planck Institute for Plant Breeding Research) for advice on the experimental method of measuring chitin-triggered ROS production in Marchantia, Ms. Yumiko Miyauchi (The University of Tokyo) for her support of gel filtration and ITC experiments, and Prof. Dr. Takayuki Kohchi (Kyoto University) for providing the wild-type male accession line Takaragaike-1 (Tak-1).

## Data availability statement

The data that support the findings of this study are available in this article or its supplementary material.

## Supporting information

Additional supporting information may be found online in the Supporting Information section at the end of the article:

Fig. S1. Conservation of EF-hand motifs among land plant RBOHs

Fig. S2. ROS-producing activity of EF-hand mutants of MpRBOHB

Fig. S3. Summary of CRISPR/Cas9-mediated genome editing of Mp*rboh* mutant lines

Fig. S4. Chitin-triggered ROS production in Mp*rboh* mutant lines

Fig. S5. Conservation of the two phosphorylation sites among land plant RBOHs Fig. S6. Estimation of intracellular Ca^2+^ concentration in HEK293T cells

Fig. S7. Preparation of recombinant proteins of the conserved regulatory domain of MpRBOHB for isothermal titration calorimetry

## Notes

### Competing Interest Statement

The authors have declared no competing interest.

