## Supporting Data for "Enhanced Ca^2+^ Binding to EF-Hands through Phosphorylation of Conserved Serine Residues Activates MpRBOHB and Chitin-Triggered ROS Production"

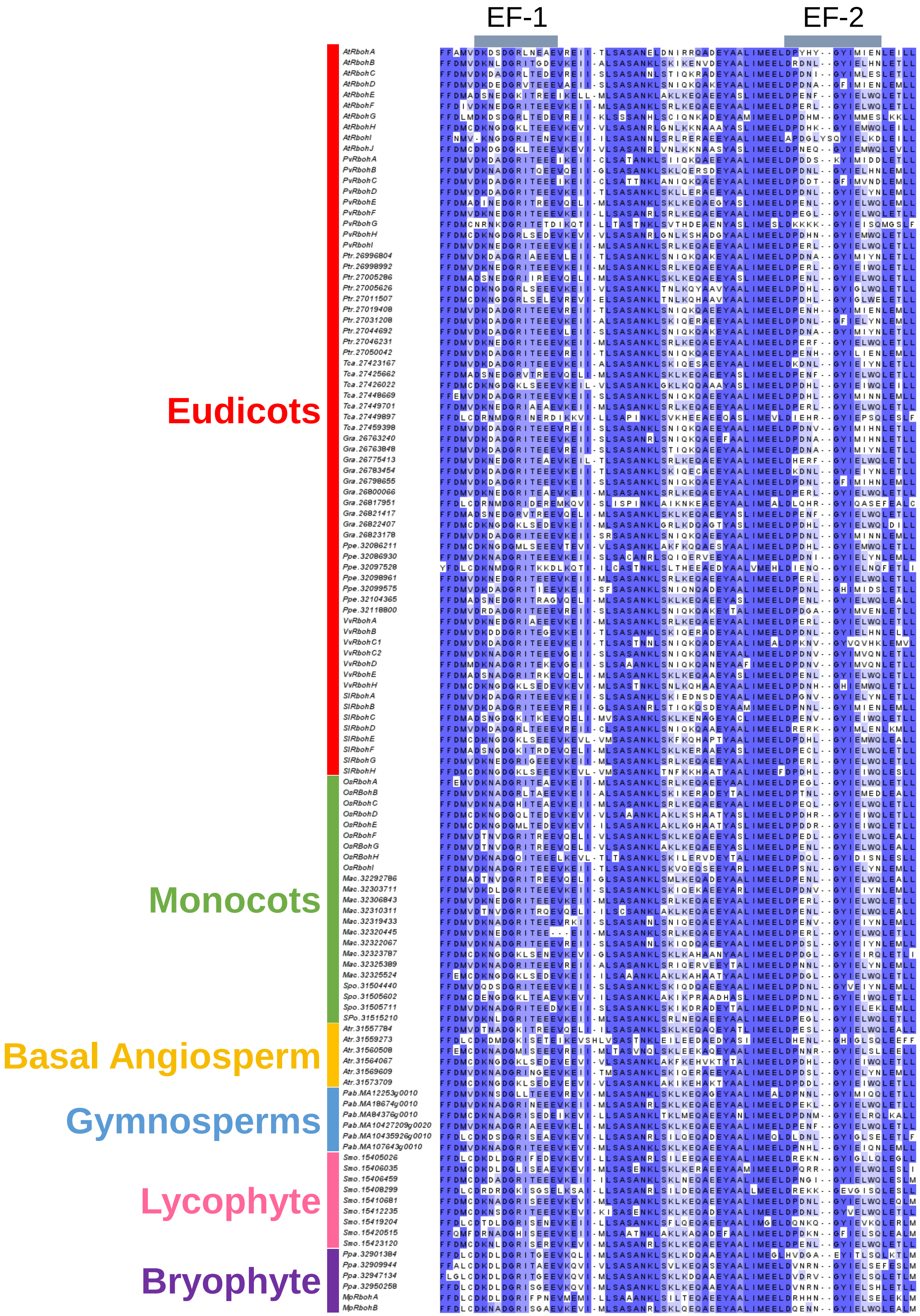


**Fig. S1. Conservation of EF-hand motifs among land plant RBOHs**

Multiple sequence alignment for 118 RBOHs from 16 plant species. It was performed by MAFFT7. Highly conserved residues are overlaid with boxes in dark blue, and relatively identical residues are in light blue. A part of the conserved N-terminal domain containing two putative $Ca^{2+}$-binding loops of EF-hand motifs is shown here, and the putative $Ca^{2+}$ binding loops are indicated by gray boxes on the alignment.

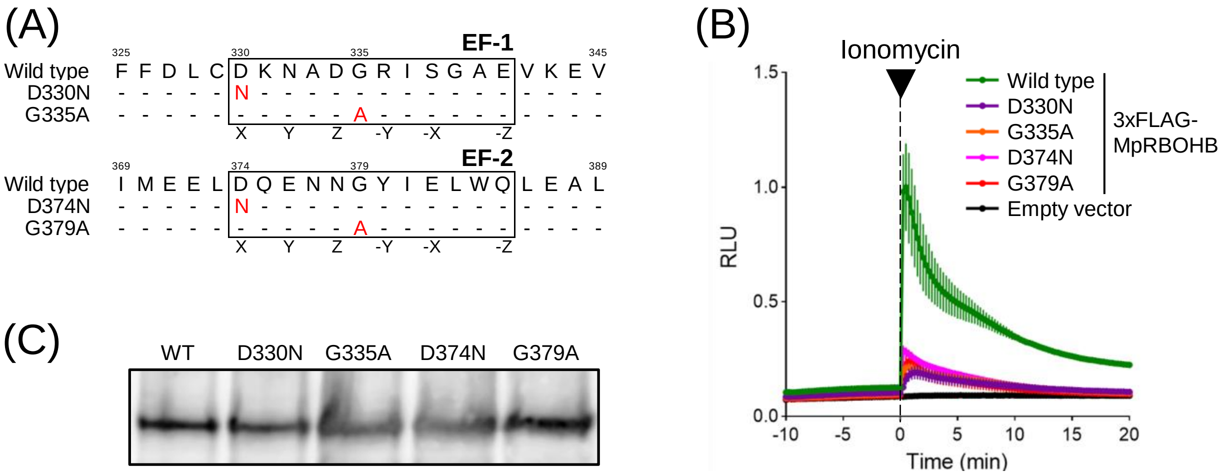


**Fig. S2. ROS-producing activity of EF-hand mutants of MpRBOHB**

(A) Amino acid sequences of the EF-hand regions of MpRBOHB. Amino acid substitutions in the MpRBOHB variants tested in this study are shown below. (B) Ionomycin-induced activation of the MpRBOHB variants in HEK293T cells. (C) Western analysis using an anti-FLAG tag antibody confirmed the expression of the MpRBOHB variants in the HEK293T cells. Data are means ± SD for three replicates.

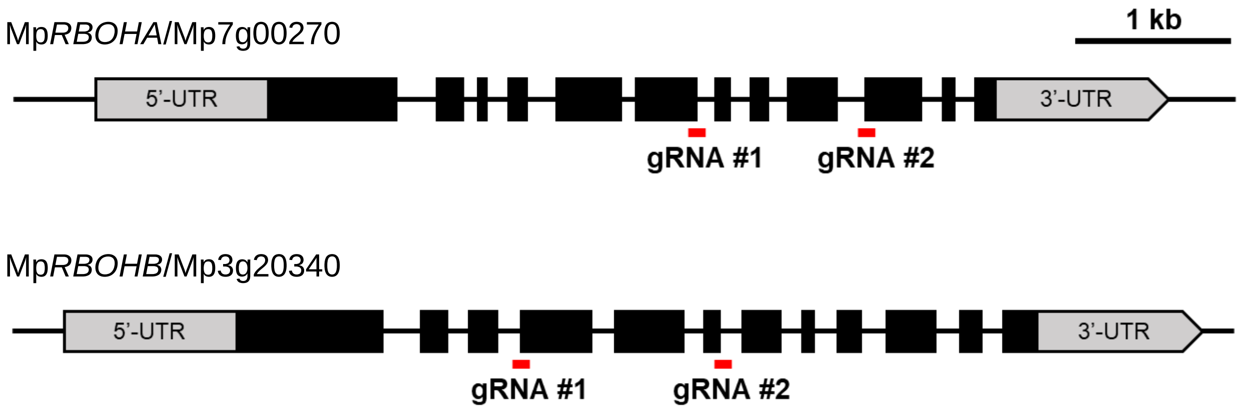


Tak-1 (male) background

| Line | gRNA target site | | Mutations confirmed by sequencing | |
|---|---|---|---|---|
| *rboha-3 ge* | #1 | PAM<br>GGTGTGTTGCGCAGGACAACAACCCCGGGGTTTG<br>intron¦exon | GGTGTG-------------------------------- | 625-base-deletion leads frame shift |
| *rboha-4 ge* | #2 | PAM<br>AGTGGAGCCAGAAGACGGTGAGCATCGGGTTCTCG<br>exon¦intron | AGTGGAGCCAGAAC-<br>CGGTGAGCATCGGGTTCTCG | 2-base-deletion and 1-base-insertion leads frame shift |
| *rbohb-1 ge* | #1 | PAM<br>TGCGAATTGGGATGTGAATATGCAGCTCTGGCAGC<br>intron¦exon | TGCGAATTGGGATGTGAATATGC<br>AGCAGCTCGAGGCCTTGATGCA<br>TATGAGCCTATGAGATATGCAGC<br>TGCATATTAGCCTATGAGTTTAG<br>GGTTGAGCTGCCTCTGGCAGC | 77-base-insertion leads frame shift |
| *rbohb-5 ge* | #2 | PAM<br>GTGACCGTTGTGAAGGTACATTCCTCAACTCCC<br>exon¦intron | GTGACCGTTG------------------------- | 766-base-deletion leads frame shift |

**Fig. S3. The summary of CRISPR/Cas9-mediated genome editing of Mp*rboh* mutant lines**

The schematic representation shows the gene structure of Mp*RBOHA* and Mp*RBOHB* and the positions of the gRNA target sites used in this study. Mp*rbohA-3ge*, Mp*rbohA-4ge*, Mp*rbohB-1ge*, and Mp*rbohB-5ge* were generated based on the male wild type (Tak-1). In the tables, underlined nucleotide sequences indicate the gRNA target sites, and the PAM sequences are highlighted in red. Mutations confirmed by sequencing the corresponding genome region are shown on the right of the tables.

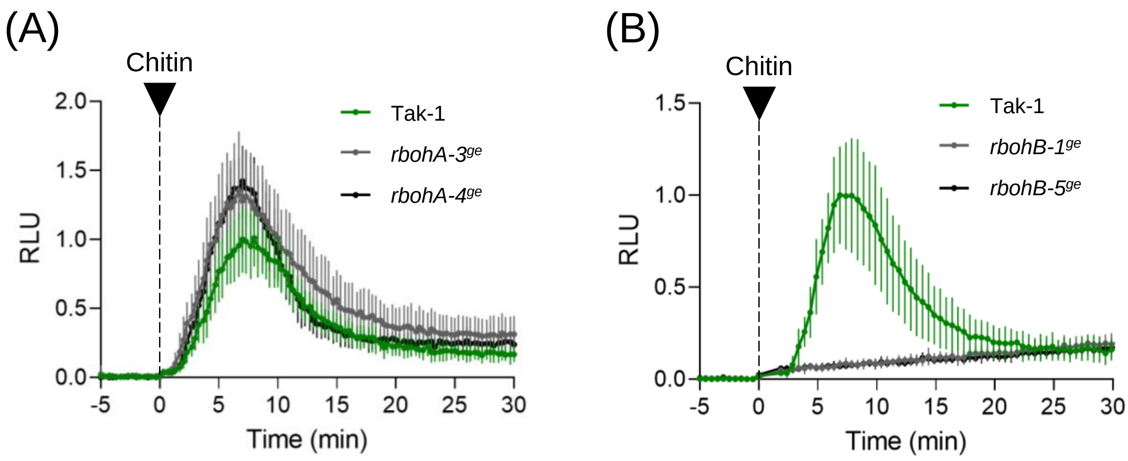


**Fig. S4. Chitin-triggered ROS production in Mp*rboh* mutant lines**

Chitin oligosaccharides-induced ROS production was measured by L-012 chemiluminescence in 7-day-old gemmalings of (A) Mp*rbohA* and (B) Mp*rbohB* mutant lines. Data are means ± SD for six replicates.

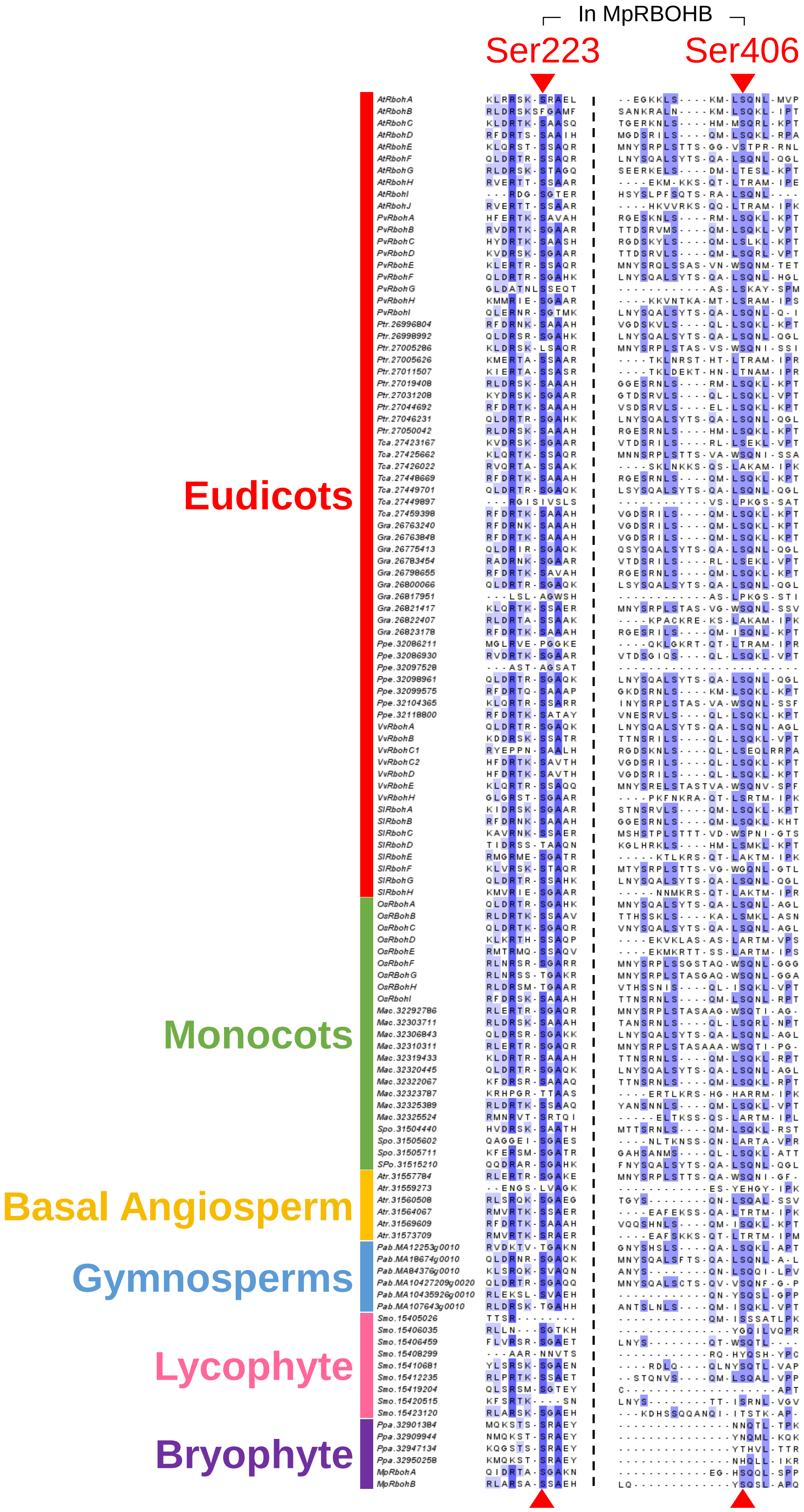


**Fig. S5. Conservation of the two phosphorylation sites among land plant RBOHs**
Multiple sequence alignment for 118 RBOHs from 16 plant species. It was performed by MAFFT7. Highly conserved residues are overlaid with boxes in dark blue, and relatively identical residues are in light blue. A part of the conserved N-terminal regions containing two Ser residues focused on in this study are shown here, and the Ser residues S223 and S406 in MpRBOHB are indicated by red arrowheads.

(A)

| $[Ca^{2+}]$ (μM) | $F^{340}$ | $F^{380}$ | Ratio ($F^{340}/F^{380}$) |
|---|---|---|---|
| 0 | 15289 | 43240 | 0.35358 |
| 0.012 | 19542 | 42957 | 0.45492 |
| 0.027 | 22391 | 40993 | 0.54622 |
| 0.046 | 24461 | 38355 | 0.63775 |
| 0.072 | 26815 | 35556 | 0.75416 |
| 0.108 | 29058 | 32784 | 0.88635 |
| 0.162 | 30920 | 29072 | 1.0636 |
| 0.252 | 32205 | 24919 | 1.2924 |
| 0.432 | 34316 | 20286 | 1.6916 |
| 0.971 | 36594 | 14396 | 2.5420 |
| 39 | 37307 | 6963 | 5.358 |

(B)

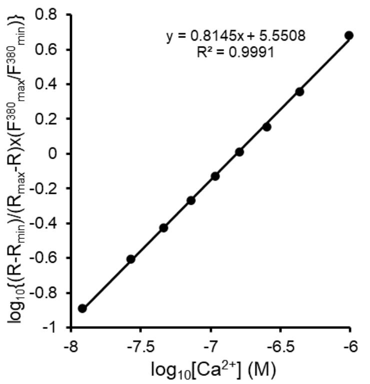


(C)

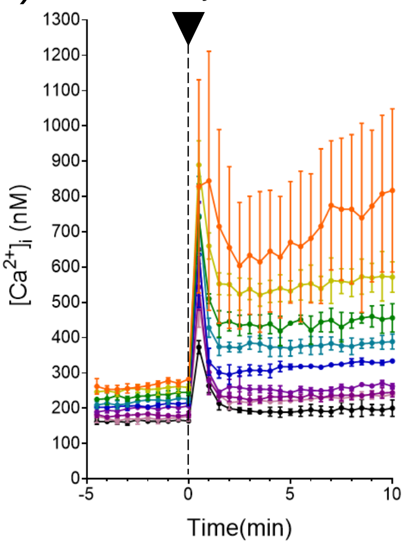


| $[Ca^{2+}]_{ex}$ (μM) | $[Ca^{2+}]_i$ (nM) | $\log_{10}[Ca]_i$ (M) |
|---|---|---|
| 2000 | 675.6 | -6.17 |
| 1000 | 567.0 | -6.25 |
| 500 | 459.6 | -6.34 |
| 250 | 399.4 | -6.40 |
| 125 | 326.6 | -6.49 |
| 62.5 | 277.6 | -6.56 |
| 31.3 | 251.2 | -6.60 |
| 15.6 | 243.6 | -6.61 |
| 0 | 210.6 | -6.68 |

**Fig. S6. Estimation of intracellular $Ca^{2+}$ concentration in HEK293T cells**

The intracellular $Ca^{2+}$ concentration ($[Ca^{2+}]_i$) in HEK293T cells was estimated by measuring Fura 2-AM fluorescence. (A, B) Fura 2 fluorescence under 0 μM to 39 μM free $Ca^{2+}$ solution was determined at 37 ° C using a micro-plate reader operating at an emission wavelength of 510 nm and excitation wavelength of 340 nm and 380 nm, and the calibration curve was plotted. $F^{340}$: a fluorescence excited by a wavelength of 340 nm, $F^{380}$: a fluorescence excited by a wavelength of 380 nm, R: $F^{340}/F^{380}$ ratio, $R_{min}$: R under 0 μM free $Ca^{2+}$, $R_{max}$: R under 39 μM free $Ca^{2+}$. (C) $F^{340}$ and $F^{380}$ of Fura 2-AM loaded HEK293T cells were measured under HBSS (-/-) buffer added 0 mM to 2 mM $CaCl_2$. 1 mM ionomycin was added at the time point indicated by the arrowhead. The left panel indicates $[Ca^{2+}]_i$ changes calculated based on the calibration curve. Data are means ± SD for three replicates. The right panel indicates the means of $[Ca^{2+}]_i$ for 5 min after ionomycin treatment under each extracellular $Ca^{2+}$ conditions.

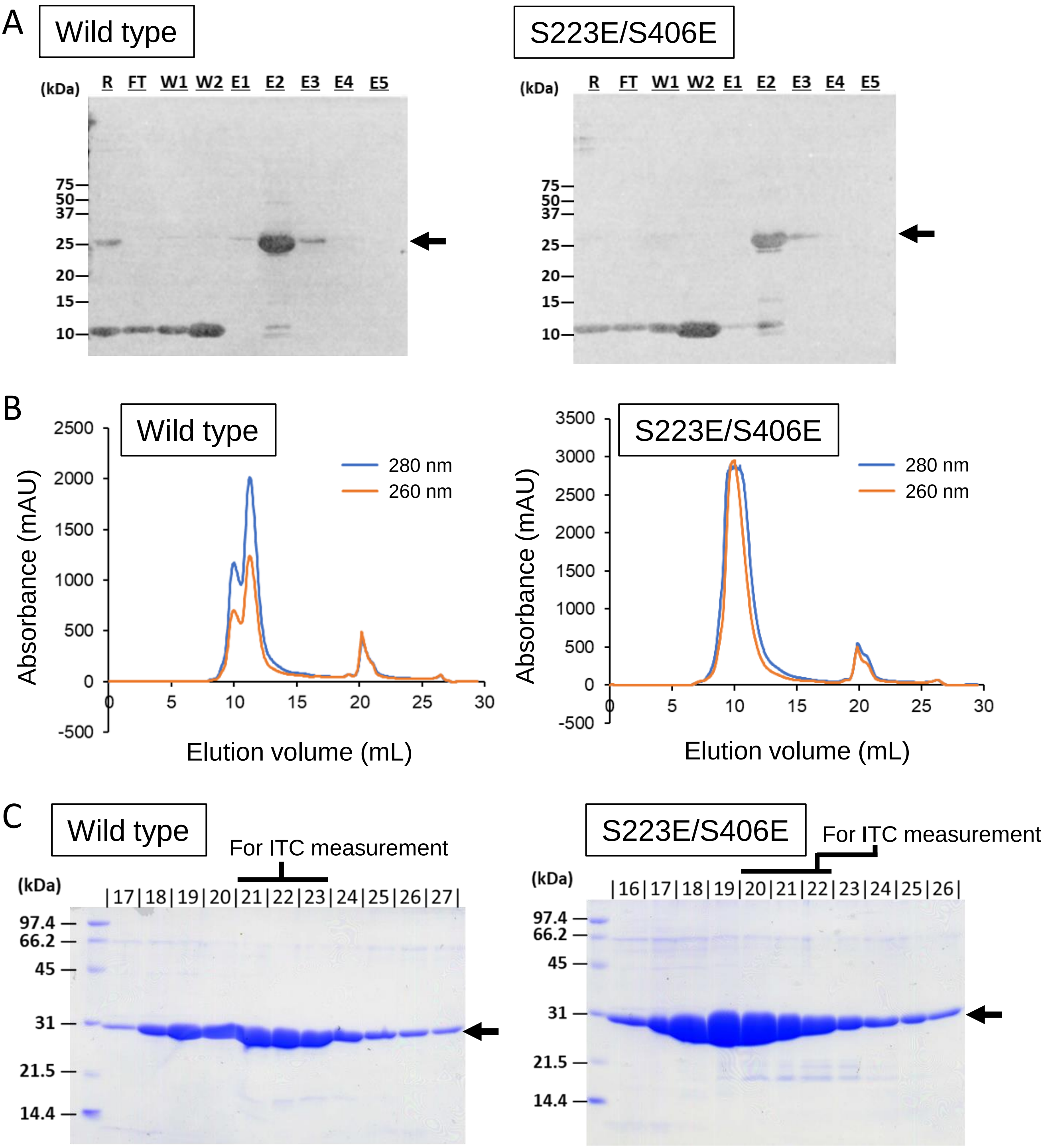


**Fig. S7. Preparation of recombinant proteins of the conserved regulatory domain in MpRBOHB for isothermal titration calorimetry**

6xHis-tagged MpRBOHB-(212-414) (MpRBOHB$^{N}$) recombinant proteins were purified for isothermal titration calorimetry (ITC) measurements. (A) Western analysis using an anti-6xHis tag antibody confirmed the MpRBOHB$^{N}$ in each fraction of primary purification by nickel-affinity chromatography. Arrows indicate MpRBOHB$^{N}$. R: raw extraction, FT: flow through, W: wash, E: elution. (B) Chromatograms of secondary purification by gel filtration were shown. These indicate absorbances at 260 nm, and 280 nm. (C) Gel filtrated fractions were separated by SDS-PAGE using 15% polyacrylamide gel and stained by CBB. Arrows indicate MpRBOHBN. Fractions with a line above the fraction numbers were used for measurements shown in Fig. 6.

### Table S1. Primers used in this study.

| | | |
|---|---|---|
| gRNA_MpRBOHA1_F | ctcgGTTGCGCAGGACAACAACCC | Guide RNA |
| gRNA_MpRBOHA1_R | aaacGGGTTGTTGTCCTGCGCAAC | Guide RNA |
| gRNA_MpRBOHA2_F | ctcgACCCGATGCTCACCGTCTTC | Guide RNA |
| gRNA_MpRBOHA2_R | aaacGAAGACGGTGAGCATCGGGT | Guide RNA |
| gRNA_MpRBOHB1_F | ctcgGGGATGTGAATATGCAGCTC | Guide RNA |
| gRNA_MpRBOHB1_R | aaacGAGCTGCATATTCACATCCC | Guide RNA |
| gRNA_MpRBOHB2_F | ctcgGAGGAATGTACCTTCACAA | Guide RNA |
| gRNA_MpRBOHB2_R | aaacTTGTGAAGGTACATTCCTC | Guide RNA |
| proMpRBOHB_OE_F | CTGCTTGCGTTTTTCTCTCC | Amplifying fragments for overlap extension with cDNA |
| proMpRBOHB_OE_R | CTGAGCTGTTCAACTCTC | Amplifying fragments for overlap extension with cDNA |
| MpRBOHBcDNA_OE_F | AAGCTCGTCATGAATCGATCTGG | Amplifying fragments for overlap extension with promoter |
| MpRBOHBcDNA_OE_R | ttttGGCGCGCCCTAGAAGTTTTCCTTGTGGAAC | Amplifying fragments for overlap extension with promoter |
| MpRBOHB_D330N_F | CCAGGATGCAGATCTTTTTTGATCTGTGTaACAAGAATGCAGATG | Site-directed mutagenesis |
| MpRBOHB_D330N_R | CATCTGCATTCTTGTtACACAGATCAAAAAAGATCTGCATCCTGG | Site-directed mutagenesis |
| MpRBOHB_G335A_F | CAAGAATGCAGATGCGCGCATCTCTGGGG | Site-directed mutagenesis |
| MpRBOHB_G335A_R | CCCCAGAGATGCGCGCATCTGCATTCTTG | Site-directed mutagenesis |
| MpRBOHB_D374N_F | TGATCATGGAAGAGCTCaACCAAGAGAACAATGGG | Site-directed mutagenesis |
| MpRBOHB_D374N_R | CCCATTGTTCTCTTGGTtGAGCTCTTCCATGATCA | Site-directed mutagenesis |
| MpRBOHB_G379A_F | CGACCAAGAGAACAATGCGTACATTGAGCTCTGGC | Site-directed mutagenesis |
| MpRBOHB_G379A_R | GCCAGAGCTCAATGTACGCATTGTTCTCTTGGTCG | Site-directed mutagenesis |
| MpRBOHB_S223E_F | GCTAGCGAGGTCTGCGgagTCTGCAGAGCATGCCT | Site-directed mutagenesis |
| MpRBOHB_S223E_R | AGGCATGCTCTGCAGActcCGCAGACCTCGCTAGC | Site-directed mutagenesis |
| MpRBOHB_S406E_F | AGATGCGTACTTGCAGTACgagCAGTCTTTGGCTCCACAGC | Site-directed mutagenesis |
| MpRBOHB_S406E_R | GCTGTGGAGCCAAAGACTGctcGTACTGCAAGTACGCATCT | Site-directed mutagenesis |
| MpRBOHB_S221/223/224A_F | CGGCTAGCGAGGgCTGCGgCCgCTGCAGAGCATG | Site-directed mutagenesis |
| MpRBOHB_S221/223/224A_R | CATGCTCTGCAGcGGcCGCAGcCCTCGCTAGCCG | Site-directed mutagenesis |
| MpRBOHB_S406/408A_F | GCGTACTTGCAGTACgcCCAGgCTTTGGCTCCACAGC | Site-directed mutagenesis |
| MpRBOHB_S406/408A_R | GCTGTGGAGCCAAAGcCTGGgcGTACTGCAAGTACGC | Site-directed mutagenesis |
| proMpRBOHB_F | ttttGCGGCCGCCCCTTGCCAAACAAAGTGATG | Amplifying the 5-kb upstream initiation codon |
| proMpRBOHB_R | ttttGGCGCGCCCACCCTTGACGAGCTTACAGTAGCTCC | Amplifying the 5-kb upstream initiation codon |
| In-Fusion_MpRBOHBN_F | gacgacgacaagatgagcgaaaacctgtattttcagagtAATCCCGCCAATCCTCG | Amplifying MpRBOHB-(212-414) with overlap sequences |
| In-Fusion_MpRBOHBN_R | gaggagaagcccggttaTTTGTTCCGCTGTGGAGC | Amplifying MpRBOHB-(212-414) with overlap sequences |
| In-Fusion_vector_F | GCTCATCTTGTCGTCGTC | Amplifying MpRBOHB-(212-414) with overlap sequences |
| In-Fusion_vector_R | TAACCGGGCTTCTCCTC | Amplifying MpRBOHB-(212-414) with overlap sequences |
| MpRBOHBcDNA_pcDNA_F | ttttGCGGCCGCCTAGAAGTTTTCCTTGTGGAAC | cDNA-subcloning into pcDNA3.1(-) |
| MpRBOHBcDNA_pcDNA_R | ttttTCTAGAAATCGATCTGGGGAGCTG | cDNA-subcloning into pcDNA3.1(-) |
| MpRBOHBcDNA_pENTR_F | caccATGAATCGATCTGGGGAGCTG | cDNA-subcloning into pENTR_D-TOPO |
| MpRBOHBcDNA_pENTR_R | CTAGAAGTTTTCCTTGTGGAACTCAAACC | cDNA-subcloning into pENTR_D-TOPO |
| GCaMP_F | ttttGCGGCCGCcATGGTCGACTCATCACGTCG | Subcloning into pENTR_D-TOPO |
| GCaMP_R | ttttGGCGCGCCTTACTTCGCTGTCATCATTTGTAC | Subcloning into pENTR_D-TOPO |
